# Electrical Signalling in Three Dimensional Bacterial Biofilms Using an Agent Based Fire-Diffuse-Fire Model

**DOI:** 10.1101/2023.11.17.567515

**Authors:** Victor Carneiro da Cunha Martorelli, Emmanuel Akabuogu, Rok Krašovec, Ian S. Roberts, Thomas A. Waigh

## Abstract

Agent based models were used to describe electrical signalling in bacterial biofilms in three dimensions. Specifically, wavefronts of potassium ions in *E. coli* biofilms subjected to stress from blue light were modelled from experimental data. Electrical signalling only occurs when the biofilms grow beyond a threshold size, which we have shown to vary with the *K*^+^ ion diffusivity and the *K*^+^ ion threshold concentration which triggered firing in the *fire-diffuse-fire model*. The transport of the propagating wavefronts shows super-diffusive scaling on time. *K*^+^ ion diffusivity is the main factor that affects the wavefront velocity. The *K*^+^ ion diffusivity and the firing threshold also affect the anomalous exponent for the propagation of the wavefront determining whether the wavefront is sub-diffusive or super-diffusive. The geometry of the biofilm and its relation to the mean square displacement (MSD) of the wavefront as a function of time was investigated for spherical, cylindrical, cubical and mushroom-like structures. The MSD varied significantly with geometry; an additional regime to the kinetics occurred when the potassium wavefront leaves the biofilm. Adding cylindrical defects to the biofilm, which are known to occur in *E. coli* biofilms, the wavefront MSD also had an extra kinetic regime for the propagation through the defect.

## 1 INTRODUCTION

Bacteria are the most common form of life on planet Earth. There are estimated to be 10^30^ bacteria on our planet that are divided among 10^12^ different species [1] [2]. Most bacteria spend most of their time adsorbed to surfaces in communities called biofilms, since there are many advantages for this lifestyle choice.[3]. Bacterial biofilms consist of a coat of extracellular polymeric substances (EPS) that forms around the communities of bacteria. Bacteria will communicate amongst themselves before the formation of a biofilm and the process is called *quorum sensing* e.g. they will vote on the suitability of a surface to grow a biofilm by releasing quorum auto-inducer molecules [4]. Bacteria in biofilms are 2-3 orders of magnitude more resistant to antibiotics than planktonic forms [5]. Thus the study of biofilms and how the bacteria communicate to form biofilms, can play a critical role in fighting medically relevant infections.

Electrical signalling via small charged ions (e.g. *K*^+^) is a newly discovered phenomenon that occurs in addition to quorum sensing mechanisms and allows bacteria to communicate in small clusters of bacteria and in biofilms [6]. Electrical signalling is possible since the bacteria act as electrically excitable cells (biofilms are an example of excitable matter) similar to neuronal, sensory and cardiac cells [7] [8]. Originally discovered in *B. subtilis* biofilms that experience nutrient starvation [9], similar electrical signalling phenomena have been observed in *E. coli* biofilms due to stress caused by exposure to blue light (via the creation of reactive oxygen species)[6]. Wave pulses of hyperpolarization have also been observed to propagate through biofilms of *N. gonorrhoeae* [10]. *P. aeruginosa* have also been observed to hyperpolarize in response to blue light [11].

The process of electrical signalling in biofilms is an emergent phenomenon due to the interaction of thousands of bacteria and thus lends itself to agent based modelling (ABM) [12]. Modelling the complex nature of electrical signalling in bacteria is important to understand fundamental aspects of bacterial physiology and to motivate ways to disrupt biofilms e.g. conducting materials in wound dressings are found to be effective against bacteria [13], but the biophysical mechanisms behind this activity are not well understood.

ABM involves simulations where the agents are constrained to follow a set of rules that determine their interactions and the rules can be stochastic in nature [14]. ABM can describe complex systems in terms of emergent phenomena from the constituent agents [15]. This is useful to describe bacterial communities as their behaviour depends on subtle interactions between the bacteria e.g. individual bacteria can act like completely different entities at high bacterial concentrations [2]. We previously used ABM to describe electrical signalling in two dimensional *B. subtilis* biofilms [16]. Here we extend the work to three dimensions and use it to describe experiments with the more medically relevant bacterium, *E. coli* [6]. A previous study of electrophysiological ABM in 3D was used to understand the species independent attraction of bacteria to biofilms [17]. Our study provides a detailed minimal model for electrochemical excitation of biofilms in which non-linear aspects of the action potentials are neglected. It is found to be sufficient to describe recent fluorescence microscopy experiments from our group with *E. coli* [6], although additional experiments indicate Hodgkin-Huxley modelling of the ion channel dynamics is required to describe electrical impedance spectroscopy experiments and electrical stimulation experiments with fluorescence microscopy. This will be covered in future publications. The current minimal model allows much of the physics to be handled in a simple transparent manner and provides a useful first step for modelling biofilm electrophysiology.

The new simulations allow us to explore the effect of biofilm morphology on the propagation of electrical waves, providing predictions for future experiments. Furthermore, the effect of defects can be studied (cylindrical pores are known to occur in *E. coli* biofilms [18]) and again could motivate future experiments. Finally the anomalous motion of the wavefronts can be studied [19] and the effect of potassium release kinetics quantified. The simulations provide a novel system to explore the *anomalous transport of wavefronts* in reaction-diffusion systems, which is expected to be a common but rarely explored biophysical phenomenon [20] and the simulations provide an example of emergent anomalous wavefronts from the classical diffusion of ions and the simple linear release kinetics of the ions by the bacteria.

### A. Agent based models

BSim is an open source Java software package for agent based modelling [12] that can be used to analyse complex behavioural patterns in the dynamics of cell populations [21]. An advantage of BSim is that it is a 3D platform, which makes calculations more realistic compared with previous 2D models of electrical signalling [16]. BSim provides an efficient mechanism to describe the release of potassium ions within a community of bacteria and the internal states of the agents can be carefully controlled.

BSim has two major components to describe electrical signalling in biofilms: the positions of the bacteria (the agents) and the chemical field which obeys Fick’s laws of diffusion (the *K*^+^ ions required for signalling). These main components are shown in Figure I.1 where they can interact in the system through differential equations.

**FIG. I.1.**
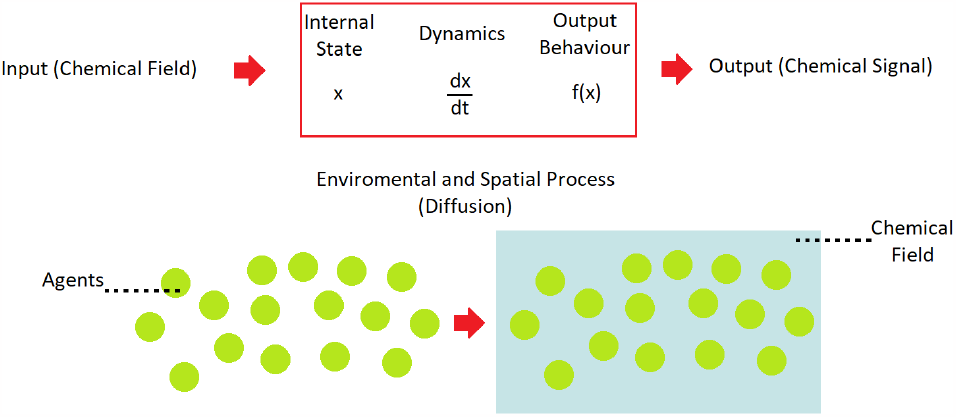
Schematic of the BSim agent based model. The two main components of the model are shown: the agents and the process by which agents interact. The agents are shown as green spheres and the chemical field is shown in blue. BSim allows the partial differential equations for the diffusion of the chemical field to be calculated. [21]

BSim was chosen for the simulation due to its versatility, which allowed the cumulative chemical field (the *K*^+^ ion concentration) from all of the bacterial agents to be calculated. Combined with the user input to determine where the bacteria are located, this allowed any biofilm shape (e.g. a spherical biofilm) to be modelled and the chemical field analysed as a function of time and space. Numerical solution to Fick’s second law of diffusion was used to calculate the transport of *K*^+^ ions for the *fire-diffuse-fire* model.

### B. Fire-Diffuse-Fire Model

A *fire-diffuse-fire* (FDF) model was used by Blee et al to describe electrical signalling in 2D bacterial biofilms [16]. It was originally developed to describe intracellular calcium signalling from discrete sites in eukaryotic cells (*Xenopus oocytes*) to model waves of propagation [7] [22] [23] [24]. Analytic solutions do not exist for the FDF model for randomly placed sources in two or three dimensions, so agent based models are needed.

In our FDF model, a spike of potassium was initiated at a specific release site, in an array of stationary bacteria that form the biofilm. When this initial spike propagates via diffusion and reaches a potassium concentration *c*\* (the firing threshold) at the position of another bacterium, an additional finite amount (*σ*) of potassium ions will be released. This will trigger a chain reaction across the whole biofilm and the spreading wave of potassium can be numerically analysed. The system resembles a wildfire in a forest.

The model uses Fick’s second law with reaction terms for the release of potassium ions from the bacteria, i.e. it is a variety of reaction-diffusion equation,

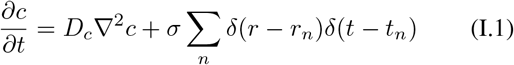

where Fick’s second law of diffusion 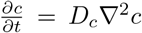 is used with *c* the *K*^+^ ion concentration, *r* is the position and *t* is the time, coupled with a reaction term for the release of potassium ions from each bacterium. Equation (I.1) considers a finite number of bacterial agents *n* which release an amount *σ* of *K*^+^ ions into the system at time, *t*_*n*_, and at the position of the bacteria, *r*_*n*_, when the *K*^+^ concentration is above the firing threshold value *c*\*.

The FDF model (I.1) predicts that the speed of the wavefront is proportional to the diffusion coefficient, which is not observed in experiments [6]. Furthermore, there should be propagation failure of the wavefronts if the diffusion coefficient (*D*_*c*_) is sufficiently small, which is not observed. This is due to the model not allowing the potassium ions to be conserved, which would be more realistic. By adding a constant decay term for the *K*^+^ ions, this problem can be solved,

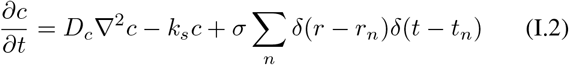

where *k*_*s*_ is a constant which allows the ions to be removed from the system e.g. due to reuptake by the bacteria. This model was studied in two dimensions by Blee *et. Al* [16] and data from fluorescence microscopy experiments closely resembles the model with *B. subtilis* biofilms grown in 2D. Although the FDF model has its limitations, it is more successful than other standard models, such as *fire and forget* in which the bacteria fire randomly [25]. As the bacteria are acting as a community, the FDF model is more satisfactory in that it can describe the synchronisation of the wavefronts[25]. Furthermore, it is possible to motivate the anomalous diffusion of wave fronts via FDF ABM.

### C. Anomalous Diffusion of Wave Fronts

It is interesting to study the dynamics of wave fronts caused by reaction-diffusion phenomena, since it will affect the timing of biological events and the patterns formed [26]. Analytic approaches have predominantly investigated diffusive or ballistic scaling based on analytic solutions to reaction-diffusion equations, but in general anomalous dynamics are possible demonstrating intermediate regimes of behaviour. Thus the propagation of wavefronts during electrical signalling in bacteria may not be as straightforward as simple Brownian diffusion, although the *K*^+^ ions are modelled using Fick’s second law in equ. (1.2). Anomalous diffusion has been observed in experiments with electrochemical wavefronts [16] [9] and it is an emergent phenomenon from simulations in 2D [16].

The mean square displacement of the wavefront position (*R*^2^) is a useful method to describe how transport occurs in a reaction-diffusion system [27]. *R*^2^ is analogous to the mean square displacement (*MSD*) of particles used to define the anomalous transport of particles [28] i.e.

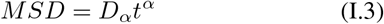

where *D*_*α*_ is the generalized diffusion coefficient and *α* is the scaling exponent. The wavefront of the *K*^+^ ion concentration field can be described using [27],

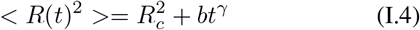

where the mean square displacement *< R*(*t*)^2^ *>* of the wavefront position is dependent on a critical radius that will initiate propagation of the wavefront (*R*_*c*_), a constant *b*, time *t* and the anomalous scaling constant *γ*. If *γ* = 1, the motion of the wavefront is defined to be *diffusive, γ <* 1, the motion is *subdiffusive*, 1 *< γ <* 2 the motion is *subballistic-superdiffusive* and if *γ >* 2 the motion is *superballistic*. Anomalous transport arises when *γ* ≠ 0, 1, 2 [29]. The critical radius (*R*_*c*_) is an emergent phenomenon from the simulations and is required to describe the experiments with *E. coli* [6].

## II. COMPUTATIONAL METHODS

The parameters for the agent based model include: the rate the *K*^+^ ions are removed from the system (*k*_*s*_), the diffusivity of the *K*^+^ ions in the biofilm (*D*_*c*_), the time step, the total simulation time, the bounds of the simulation box, the total amount of *K*^+^ ions and the threshold concentration for each bacterium to fire (*c*\*). When the threshold concentration is surpassed at the position of a bacterium it will release an additional amount of *K*^+^ (*σ*) at its position. There are also time constraints applied for the first peak and second peak of the FDF model, since two propagating wavefronts are observed in *E. coli* experiments in response to light [6], presumably due to the dynamics of a gene network inside each bacterium that handles their response to the reactive oxygen species (ROS) the light creates (a model for adaptation to ROS stress was presented [6]). The total number of bacteria is also user defined.

A spherical geometry was first used to describe the biofilm (Fig. II.1). The bacteria were static i.e. immotile and not dividing on the time scale of the experiment. The bacterial agents were arranged randomly within the workspace by defining two characteristic spherical coordinates *θ* and *ϕ*,

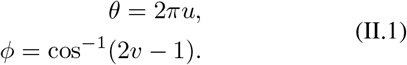

The variables *u* and *v* were defined as random numbers from 0 to 1. Therefore, random angles for *θ* and *ϕ* were obtained for the spherical coordinates. The random angles were assigned to their respective Cartesian coordinates *x*_0_, *y*_0_ and *z*_0_ using the transformation,

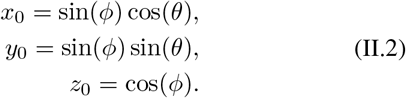

This takes a random point from 0 to 1 for each coordinate that will fall inside a unit sphere. The points in the workspace of the sphere (*x*_00_, *y*_00_, *z*_00_) are then assigned,

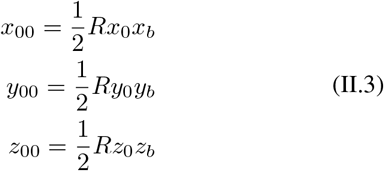

where *R* is the cube root of a random variable from 0 to 1, which will help ensure a spherical symmetry within the workspace. *x*_*b*_, *y*_*b*_ and *z*_*b*_ are the boundary extents of the workspace. A sphere is created with radius half the size of the workspace. An extra restriction ensures the bacteria do not overlap by assigning an “if” statement to delete attempts to place bacteria within *r* (the size of a bacterium) of the same location. Figure II.1 shows a spherical biofilm generated with 6000 bacteria. Both reflecting and periodic boundary conditions can be implemented for calculations of the chemical field (the *K*^+^ ion concentration).

**FIG. II.1.**
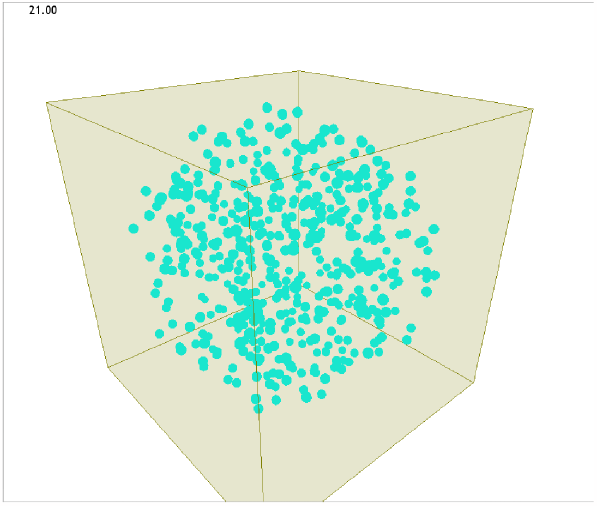
An example of a spherical biofilm constructed using BSim. The bacteria are stationary and are shown in blue. The EPS is neglected in the model. The bacteria are randomly placed within the sphere, but bacterial overlaps are not permitted. The simulation box is a cube (grey).

In all the biofilm geometries explored, the bacteria are initially spiked only once and there can be more than one bacteria in the volume of the initial spike. The result of the initial stimulation is thus dependent on the number of bacteria. Propagation failure can occur if no bacteria are found in the region of initial spiking. However, this is unlikely due to the total number of bacteria used and was not observed in the current simulations.

After the initial stimulation, the wavefront began to propagate through the biofilm. If the time of the stimulation is shorter than an initial wavefront parameter (in seconds), which is defined as the time over which the wavefront will stop propagating, the FDF mechanism continues. If the stimulation time is lower than a second wavefront parameter (in seconds), which is defined as the time at which the wavefront will begin propagating again (the second wavefront), but higher than the initial wavefront parameter, then the loss mechanism dominates i.e the fire-diffuse-fire mechanism stops and the wavefront stops propagating. Finally, if the stimulation time is higher than the second wavefront time parameter, it will continuously propagate the FDF mechanism. After all of the conditions and statements are completed, each time step is finished and the chemical field gets updated. One can toggle between showing the bacteria and the chemical field (the *K*^+^ concentration).

### A. Measuring Wavefront Velocity

The wavefront position was defined using the threshold concentration (*c*\*) of the potassium ions for propagation. The code has a loop that checks the ion concentrations first in time and then in space. When the potassium ion concentration is higher than the threshold (*c*\*), a value of 1 is assigned to that radius, so that the wavefront is known to be at that position. Afterwards, the indices of the 1s are all retrieved, which gives the time and space coordinates for the wavefront. If the wavefront is inwards rather than outwards, then simply flipping the array and assigning the wavefront position for values below the threshold provides a useful definition.

Once the position of the wavefront is found, it is straightforward to plot the time against the position and fit a polynomial. The derivative of the curve is a reasonable approximation to the instantaneous velocity of the wavefront at different points in time and space.

## III. RESULTS

### A. *K*^+^ Ion Diffusion from a Single Bacterium

To confirm that the diffusion code functions accurately in Bsim and is consistent with Fick’s laws of diffusion, the diffusion profile for release of potassium ions from a single bacterium was investigated. A single bacterium was placed in the centre of the 3D workspace and it was fired once. The diffusion profile of the *K*^+^ ions at different times was then calculated for a single bacterium as a function of distance.

In Figure III.1, Fick’s laws of diffusion appear to be simulated correctly for a single bacterium. The ion concentration follows a Gaussian form with log Intensity proportional to *r*^2^. The slight asymmetry in the logarithmic concentration profiles (Figure III.1a) are due to the discrete positioning of the bacterium magnified by the logarithmic scale i.e. a slight asymmetry exists due to a one pixel offset.

**FIG. III.1.**
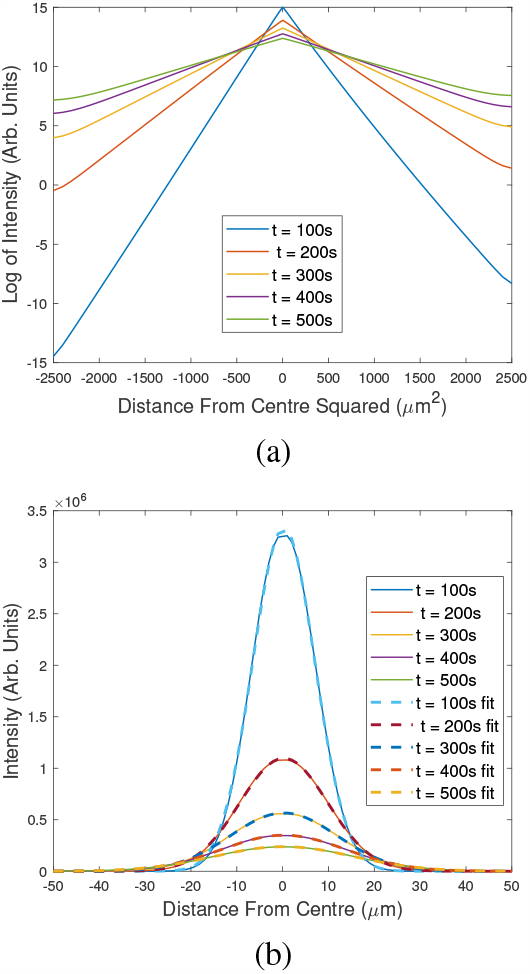
(a) Simulation of the release of potassium ions of a single bacterium over different time intervals. A single spike occurs in the middle, where the single cell was located as expected from Fick’s laws of diffusion, and a decrease in the peak heights occurs over time. A Gaussian curve of the form 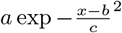 was fit for each of the time curves. For 100, 200, 300, 400 and 500 s the values for *a* were 3.33*×*10^6^, 1.10*×*10^6^, 5.66*×*10^5^, 3.50*×*10^5^ and 2.39 10^5^ respectively; the values for *b* were 2.54, 2.54, 2.55, 2.55 and 2.56 respectively; the value for *c* were 9.64, 13.6, 16.6, 19.1 and 21.3 respectively. The fits all corresponded to a *R*^2^ value of 0.9999. (b) The logarithm of the intensity and the square of the distance were considered to ensure it is Gaussian. The straight lines suggest that the trend is of Gaussian nature, though it is slightly off-centre due to the discrete nature of the lattice i.e. a one pixel bias. The potassium diffusivity (*D*_*c*_) was 0.1 *µm*^2^*/s* and the decay constant (*k*_*s*_) was set to a value of 4.44 *×* 10^*−*4^. The quantity of potassium ions added (*σ*) was 5 *×* 10^9^.

### B. Average Wavefront Velocity and *K*^+^ Ion Diffusivity

The average velocity of the centripetal (outward moving) wavefront was calculated at each point along Cartesian axes superposed on the spherical biofilm. A movie of a complete simulation is shown in the Supplementary Video in which the bacterial agents are invisible to allow a better view of the chemical field propagation. These agents however, were located in the shape shown in Figure II.1 and were present in the calculations.

Figure III.2 shows the average velocity of the *K*^+^ wavefront as a function of the diffusivity of the *K*^+^ ions. As expected, the average wavefront velocity increases as the *K*^+^ diffusivity increases. *K*^+^ ions diffuse at a constant rate in bulk water consistent with the Stokes-Einstein equation [30], but extra cellular polymeric substance in the biofilm can reduce the value of effective diffusion coefficients (more EPS will reduce *D*_*c*_) and thus it is interesting to explore the effects of this parameter. Power law fits of the average wavefront velocity on *K*^+^ ion diffusivity gives the scaling exponent *b* = 1.85.

### C. Instantaneous Velocity and *K*^+^ Ion Diffusivity

To measure the instantaneous velocity of the *K*^+^ wavefronts, a polynomial curve was fit to the position of the wavefront as a function of time. Then the time derivative was taken to give the instantaneous velocity.

**FIG. III.2.**
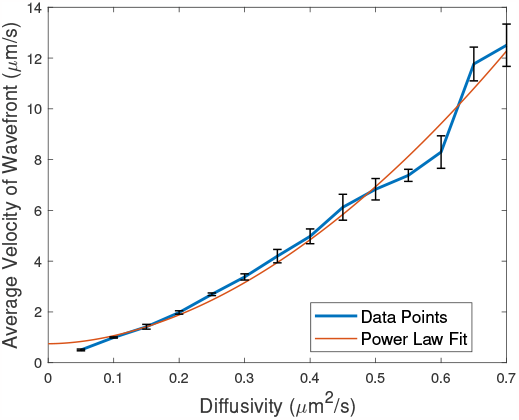
Average velocity of a wavefront of *K*^+^ ions as a function of ion diffusivity, averaged over 3 simulations. The decay rate (*k*_*s*_) was kept constant at 4.44 *×*10^*−*4^ molecules per second and the *K*^+^ ions released was kept constant at 5*×*10^9^ molecules in the FDF model for the whole biofilm. The firing threshold (c*) was maintained at 7*×*10^6^. The method for determining the intensities was the longitudinal line method in blue and spherical shell method in red. They are in good agreement. The error bars show the standard error on the mean of 3 runs. The power law fit was of the form *f* (*x*) = *ax*^*b*^ + *c*, where, *a* = 22.3, *b* = 1.85, and *c* = 0.75.

The instantaneous wavefront velocity calculated at different radii from the point of initial spiking is shown in Figure III.3 as a function of *K*^+^ ion diffusivity. There is a steady increase of the wavefront velocity with radius following a power law of the form of *v* = *aR*^*b*^ + *c*, where *a* = 0.978, *b* = 0.65 and *c* = 0.23. The scaling exponent is significantly lower for the instantaneous velocity than for the average velocity in Figure III.2. Figure III.4 shows how the scaling exponent *b* for the instantaneous velocity depends on the radius of the biofilm (rescaled to form a fractional radius on the figure).

**FIG. III.3.**
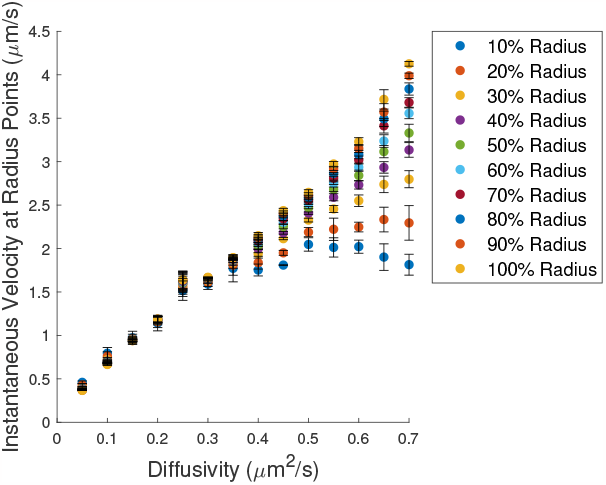
The instantaneous velocity of *K*^+^ ion wavefronts as a function of *K*^+^ diffusivity calculated for a range of radii. An average of 3 simulations is shown. The decay rate (*k*_*s*_) was kept constant at 4.44*×*10^*−*4^ molecules per second and the amount of potassium ions released (*σ*) was kept constant at 5*×*10^9^ molecules in the FDF model. The firing threshold (*c*\*) was maintained at 7*×*10^6^. The error bars are the standard error on the mean of 3 simulations. The power law fit was of the form *v*(*D*) = *aD*^*b*^ + *c*, where, *a* = 0.97, *b* = 0.65, and *c* = 0.23

**FIG. III.4.**
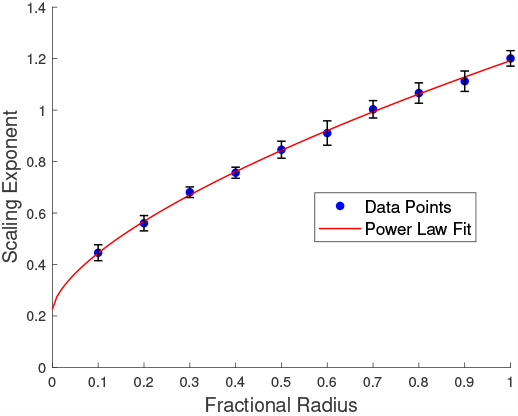
Power law scaling exponents of the instantaneous velocity of the *K*^+^ ion wavefront as a function of radial position inside the biofilm for an average of 3 runs (the fractional radius=1 on the edge of the biofilm). The decay rate (*k*_*s*_) was kept constant at 4.44*×*10^*−*4^ molecules per second and the ions released (*σ*) was kept constant at 5*×*10^9^ in the FDF model. The error bars are the standard error mean of the 3 runs.

### D. *K*^+^ Ion Profile Averaged Over the Entire Biofilm

Experimentally, two hyperpolarisation events occur in *E. coli* biofilms in response to illumination with blue light and they can be observed in globally averaged intensity profiles from fluorescence microscopy experiments using ThT as a fluorescent Nernstian probe [6]. The first hyperpolarization event is when the bacteria initially respond to the blue light and the second hyperpolarization event is thought to be a habituation phenomenon. These global oscillations could be described by the FDF ABM simulations. This is done by setting two spikes of hyperpolarisation which will trigger the FDF model and these spikes will be triggered at different time intervals which follow the experimental data. The potassium ion concentration is then globally averaged over the entire simulation box. The two hyperpolarization events are shown in Fig. III.5. The simulations are in reasonable agreement with experiment. *t*_1_ is the time for the first centripetal wavefront (the first hyperpolarisation event), *t*_2_ is the time for the centrifugal wavefront (recovery) and *t*_3_ is the time for the second centripetal event (the second hyperpolarization event). However, the second hyperpolarization event occurs more abruptly with the simulation than with the experiment.

**FIG. III.5.**
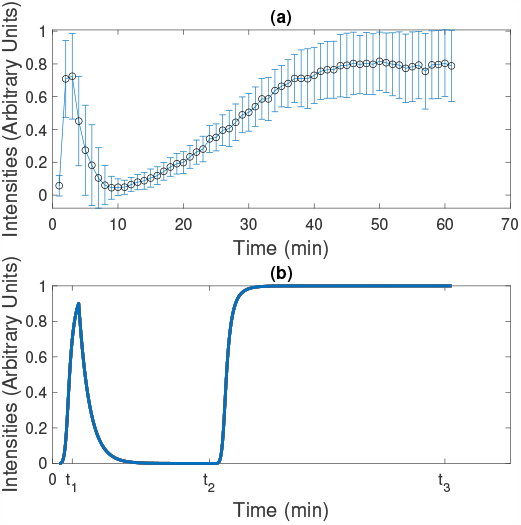
Globally averaged fluorescence intensity from a) experimental data compared to b) simulation data. a) ThT was used as a Nernstian probe in fluorescence microscopy experiments with an *E. coli* biofilm [6]. The error bars in the top subfigure represent the standard deviation over more than 3 experiments. b) The global oscillations for the bottom figure are taken for a simulation with *K*^+^ diffusivity (*D*_*c*_) of 0.1 *µm*^2^, decay rate (*k*_*c*_) of 5*×*10^*−*3^ molecules/s, 5*×*10^9^ potassium ions (*σ*) added per spike and 1*×*10^7^ potassium ions as the firing threshold (*c*\*) for the FDF mechanism. The first wavefront of the FDF stops at *t*_1_ = 180 seconds and the second wavefront begins (a new spike) at *t*_2_ = 1440 seconds. The simulation stops at *t*_3_ = 3600 seconds.

### E. Simulation of the Wavefront Dynamics

To understand the propagation of ion channel signalling in a 3D biofilm, the *K*^+^ wavefront was analysed by fitting equation I.4 to the mean square displacement of the *K*^+^ wavefront position and the equation provides an empirical definition of the *anomalous wavefront dynamics* [27]. The hyperpolarisation and depolarization wavefronts are denoted as the *centripetal* and *centrifugal* wavefronts respectively. The simulations were used to describe the experimental data.

The centrifugal *K*^+^ wavefront has a power law scaling exponent *γ* of 1.21 and is thus super-diffusive. This is shown in Figure III.6 (a) and the critical radius for propagation of the wavefront was 6.2*±*1.8 *µm*. The centripetal wavefront has a value of *γ* of 2.26 and is thus super-ballistic. This is shown in Figure III.6 (b).

**FIG. III.6.**
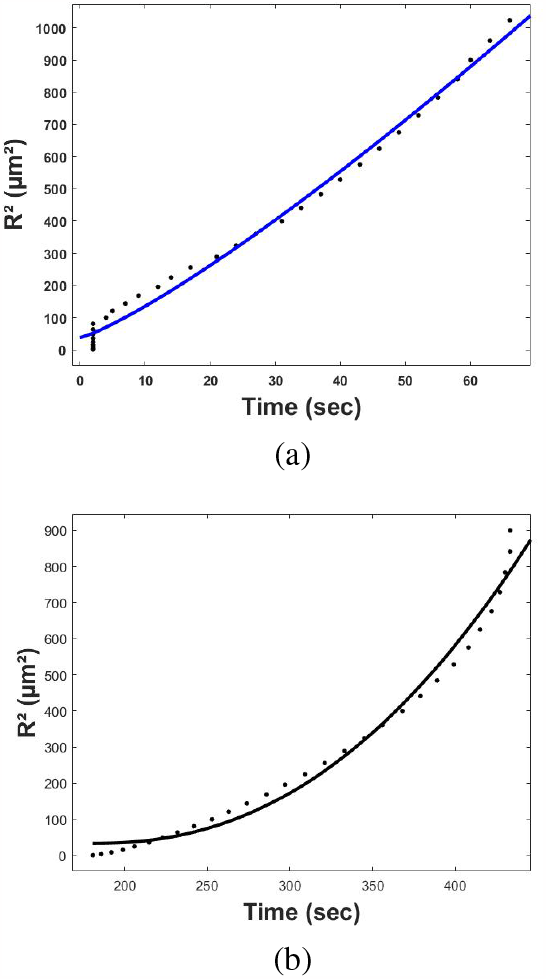
Mean square displacement of (a) centrifugal and (b) centripetal *K*^+^ wavefronts for a simulation with *K*^+^ diffusivity of 0.1 *µm*^2^, decay rate (*k*_*s*_) of 5*×*10^*−*3^ molecules/s, 5*×*10^9^ quantity of *K*^+^ (*σ*) added per hyperpolarization event and 3 *×*10^9^ molecules as the threshold (*c*\*) for the firing mechanism. (a) The curve fit is equation I.4, where *γ* is 1.21*±*0.12 and *R*_*c*_ is 6.17*±*1.84 *µm*. The error bars on the constants of the fits are 95% confidence intervals. (b) The curve fit is again equation I.4, where *γ* is 2.26*±*0.31 and *R*_*c*_ is undefined due to the nature of the centripetal wavefront. The error bars on the fits are 95% confidence intervals.

The *K*^+^ wavefront velocity as a function of time is shown in Figure III.7. The velocity of the centrifugal wavefront was almost constant with time, whereas the centripetal wavefront has a linear decrease.

**FIG. III.7.**
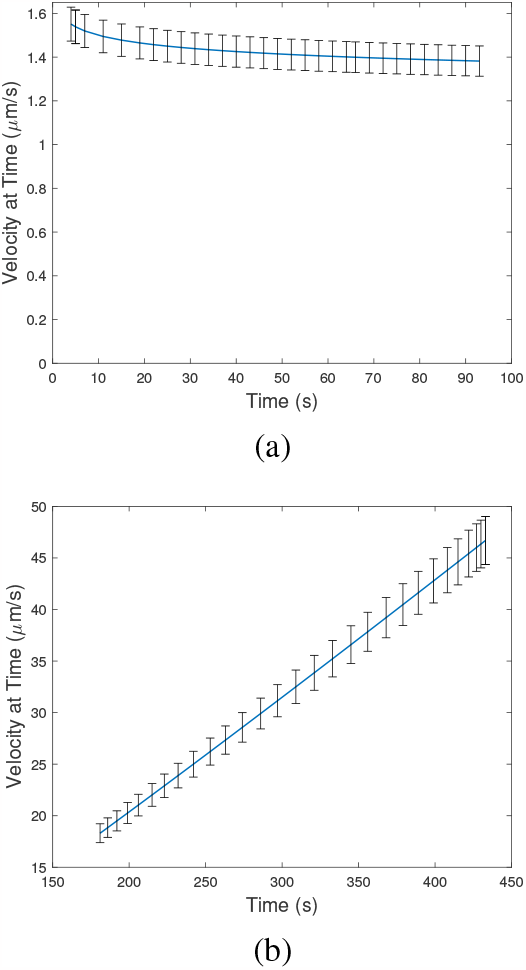
Instantaneous velocity as a function of time of (a) the centrifugal and (b) centripetal *K*^+^ wavefronts for simulations where the *K*^+^ diffusivity (*D*_*c*_) is 0.1 *µm*^2^, the decay rate (*k*_*c*_) is 5*×*10^*−*3^ molecules/s, the quantity of *K*^+^ added per hyperpolarization event (*σ*) is 5*×*10^9^ and 3*×*10^9^ molecules is the firing threshold (*c*\*) for the FDF mechanism.

Another characteristic of the *K*^+^ wavefront is how its velocity depends on its radius of curvature. In cardiac tissue (another classic example of excitable matter) a complex interplay between wavefront curvature and velocity is observed and it is related to cardiac physiology e.g. pacing of the heart with scroll waves [7]. For a *K*^+^ ion wavefront in a spherical biofilm, the curvature can be approximated by the reciprocal of the radius of the wavefront, i.e. 1*/R*.

Figure III.8 (a) shows how the instantaneous velocity of the *K*^+^ wavefront changes as a function of its curvature. The centrifugal *K*^+^ wavefront has almost no dependence on curvature, whereas the centripetal wavefront has a monotonic increase which plateaus at high curvatures [7]. Analytically the Eikonal approximation predicts the instantaneous wavefront velocity (*v*) is proportional to the curvature (1*/R*),*∝ v* 1*/R*, in reasonable agreement with the behaviour of the centrifugal wavefront [7].

**FIG. III.8.**
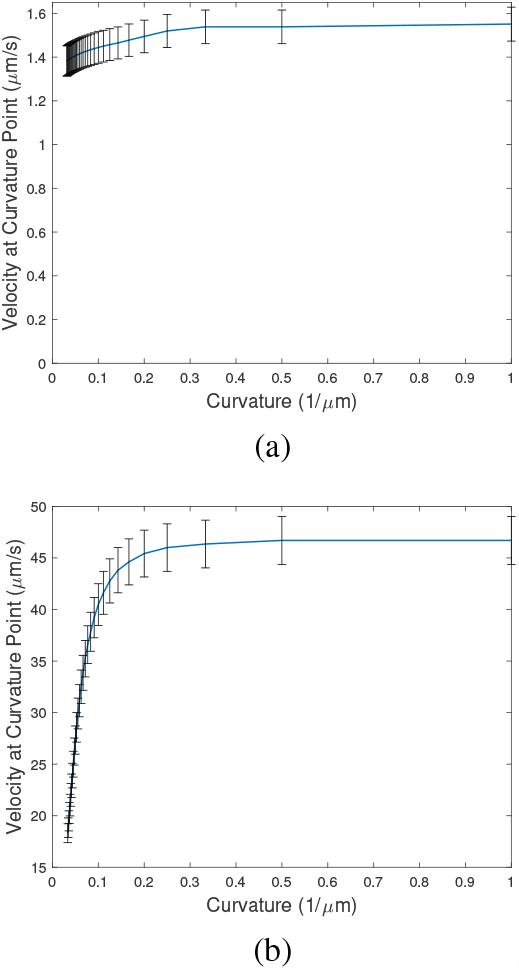
Instantaneous velocity as a function of curvature (= 1*/R*) for (a) the centrifugal and (b) the centripetal wavefronts from simulations with *K*^+^ diffusivity (*D*_*c*_) of 0.1 *µm*^2^, decay rate (*k*_*s*_) of 5*×*10^*−*3^ molecules/s, 5*×*10^9^ quantity of *K*^+^ (*σ*) added per hyperpolarization event and 3*×*10^9^ molecules as the firing threshold (*c*\*) for the FDF mechanism. The radius is defined as zero at the centre of the biofilm and the biofilm periphery is the maximum radius. The error bars show the 95% confidence interval for the velocities.

### F. Experimental Wavefront Dynamics

Confocal microscopy was used to measure the *K*^+^ wavefront dynamics in 3D using the fluorescent dye ThT as a Nernstian voltage probe. The confocal microscopy technique allows good resolution along the z-axis (transverse axis) of thick biological samples and allows vertical optical sectioning. The thickness of the biofilm shown in Figure III.9 was 154 *µm*. The sample was exposed to blue light stimulation for 60 minutes [6]. The intensity profile from confocal microscopy suggests that the biofilm begins to spike at a depth of 78 *µm*. Multiple optical sections were made at different equally spaced distances from the centre of the biofilm, perpendicular to the z-axis of the wavefront. The radial distance was measured from the core to the periphery. Figure III.9 shows the mean square displacement of the wavefront position *R*^2^ as a function of time. The critical radii and the scaling exponents *γ* are very close to the model predictions. The centrifugal wavefront anomalous exponents were 1.21*±*0.12 and 1.22*±* 0.15 for the model and experiment respectively. The critical radius was in reasonable agreement at 6.2*±*1.8 *µm* and 4.7*±* 1.0 *µm* for the model and experiment respectively.

**FIG. III.9.**
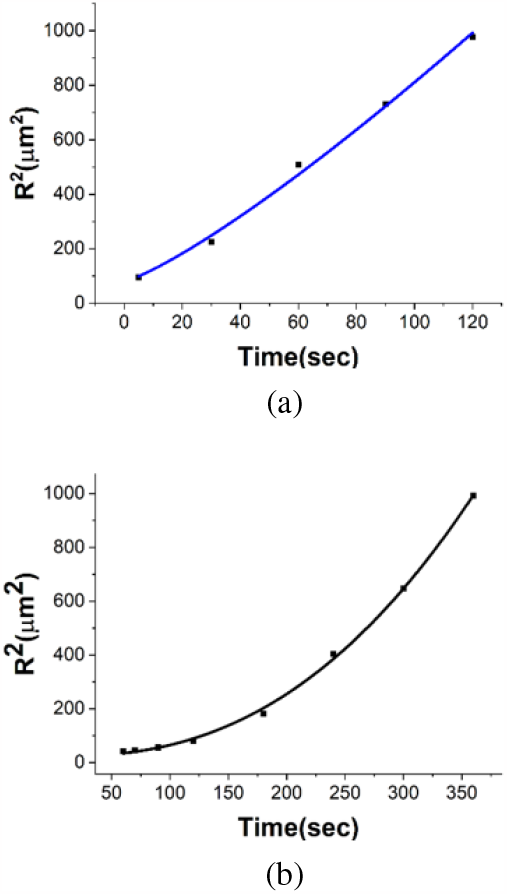
Mean square displacement of (a) the centrifugal and (b) centripetal *K*^+^ wavefronts as a function of time measured using confocal microscopy with ThT fluorophores. (a) The curve fit is equation I.4, where *γ* is 1.22*±*0.15 and *R*_*c*_ is 4.71*±*0.98 *µm*. (b) The curve fit is equation I.4, where *γ* is 2.43 0.08 and *R*_*c*_ is undefined due to the nature of the centripetal wavefront.

How the velocity varies with time and radius for the experimental confocal microscopy results was also investigated. There is a non-linear dependence of the wavefront velocity on the radius, Figure III.10, and on time, Figure III.11, for both centrifugal and centripetal wavefronts.

**FIG. III.10.**
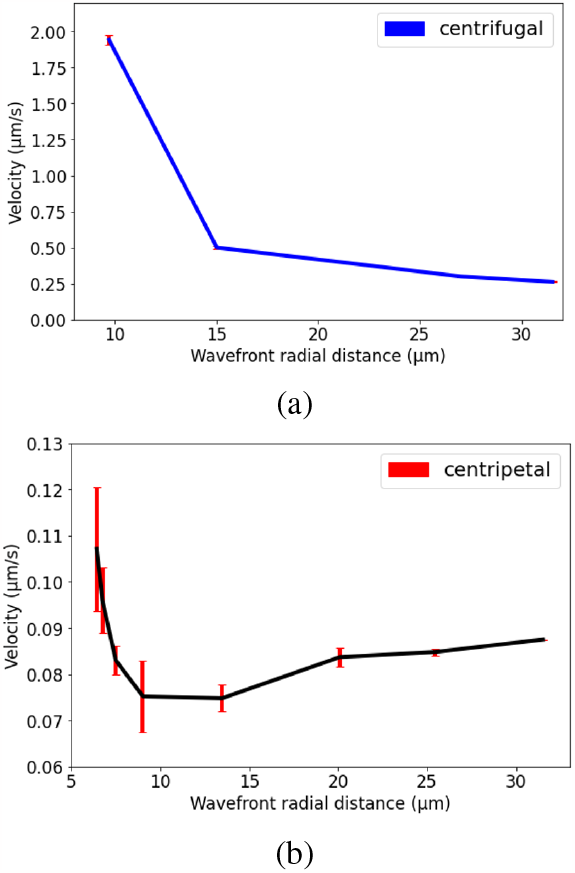
Instantaneous velocity as a function of radius of (a) the centrifugal and (b) the centripetal *K*^+^ wavefronts measured from the first peak of the confocal microscopy data. An inverse relation of the *K*^+^ wavefront velocity is observed with radius for (a), whereas (b) has a minimum. The error bars are the standard deviation of more than 3 experiments.

**FIG. III.11.**
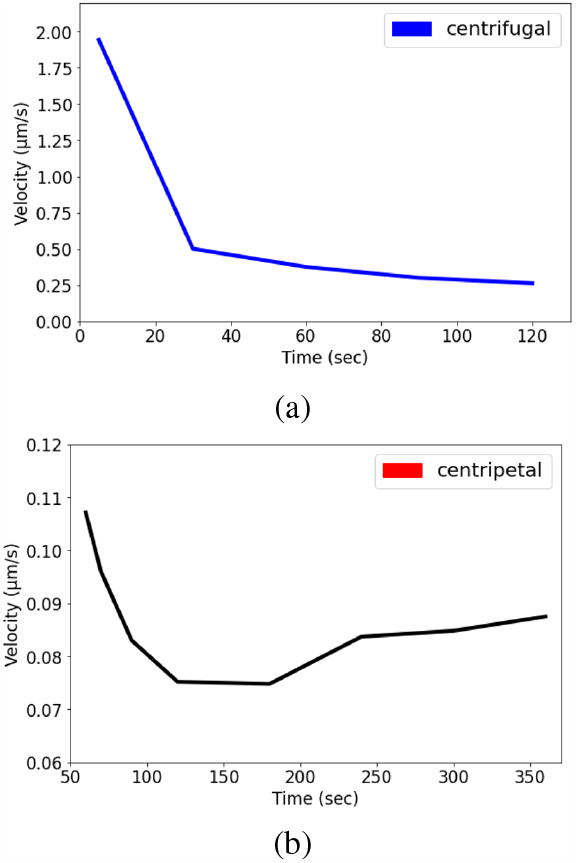
Instantaneous velocity as a function of time for (a) the centrifugal and (b) the centripetal *K*^+^ wavefronts measured from the first hyperpolarization event of the confocal microscopy data. An inverse relationship of the wavefront velocity with time is shown for (a), whereas (b) has a minimum. The error bars are the standard deviation of more than 3 experiments.

### G. Anomalous Diffusion of the Wavefront

Factors affecting anomalous diffusion of the *K*^+^ wavefronts were explored. In Figure III.12 for lower diffusivities of the potassium ions the mean square displacement position (*R*^2^) behaves in a more ballistic manner (*R*^2^*∼t*^2^), whereas for higher diffusivities it becomes more diffusive (*R*^2^*∼t*^1^).

**FIG. III.12.**
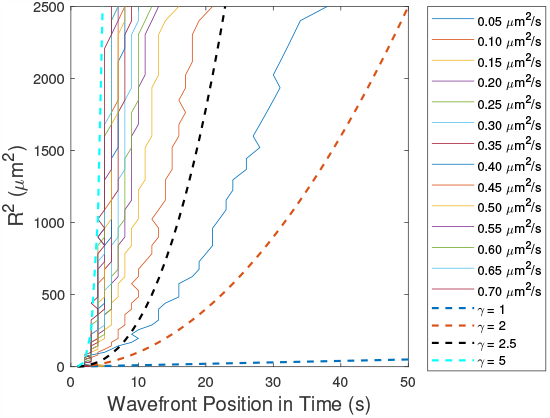
Mean square displacement of *K*^+^ wavefronts as a function of time for different *K*^+^ diffusivities from ABM simulations. The different colours represent the different *K*^+^ diffusivities shown in the legend. All *K*^+^ diffusivities are given in *µm*^2^*/s*. The dashed trendlines for different *γ* power law scaling exponents are also shown for reference.

Figure III.13 shows the mean square displacement of the *K*^+^ wavefronts as a function of time. The anomalous diffusion also varies with the firing threshold (*c*\*). This suggests that the higher the values of the threshold, the less ballistic is the wavefront i.e. the lower the anomalous scaling exponent (*α*) in equ. I.4.

**FIG. III.13.**
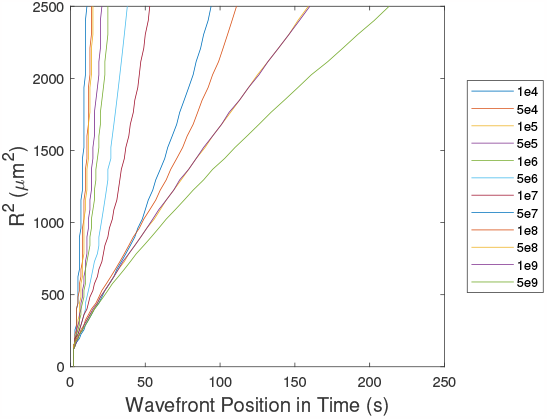
Mean square displacement of the *K*^+^ wavefronts as a function of time for different *K*^+^ concentration firing thresholds (*c*\*) from ABM simulations. The different colours represent the different firing thresholds shown in the legend. All firing thresholds are given in terms of the number of molecules needed to trigger the FDF mechanism.

### H. Critical Radius of Biofilms for Propagating Wavefronts

A critical radius for propagation of potassium wavefronts was observed in the simulations, in agreement with experiments. The critical radius is a measure of how much the initial spike needs to move before the wavefront propagates across the whole biofilm. The critical radius was found to be independent of the firing threshold (*c*\*) of the FDF model. The critical radius decreases non-linearly with the diffusivity of the potassium ions (Figure III.14) and higher *K*^+^ diffusivity creates smaller critical radii.

**FIG. III.14.**
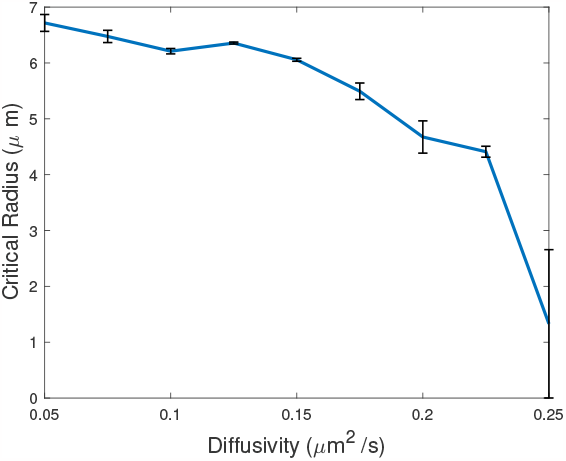
Critical radius for propagation of *K*^+^ wavefronts as a function of the diffusion coefficients of the *K*^+^ ions (*D*_*c*_, the diffusivity) from ABM simulations. An average of 3 runs is shown. The critical radii decrease as the *K*^+^ diffusivity increases. The error bars are the standard error mean of the 3 runs.

The critical radius for *K*^+^ wavefront propagation was not affected by the density of the bacteria in the biofilm in the simulations. For lower bacterial numbers, the wavefront did not propagate. Also the decay rate of potassium ions (*k*_*s*_) does not seem to influence the critical radius.

### I. Different Biofilm Geometries

Biofilms in nature grow in a variety of shapes and sizes [31] Although the spherical biofilm was useful due to its simplicity, other geometries are also interesting. Furthermore, defect structures in cylindrical biofilms were explored, since they will be common in nature e.g. foreign particles often occur in biofilms, including foreign non-spiking bacteria, and water-filled cylindrical transport pores are actively maintained by *E. coli* in their biofilms [18].

The globally averaged intensity of the hyperpolarizationevents (the *K*^+^ concentration) of the biofilm are relatively independent of the biofilm shape, Figure III.16.

**FIG. III.15.**
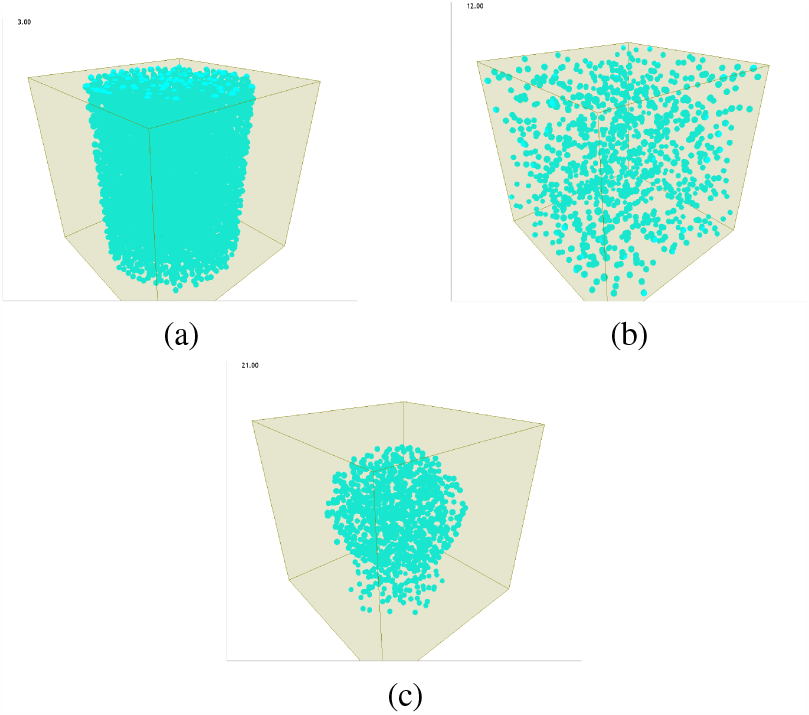
Different bacterial biofilm geometries that were used for the FDF ABM in addition to a spherical geometry (Figure II.1): (a) a cylinder, (b) a cube and (c) a mushroom shape.

**FIG. III.16.**
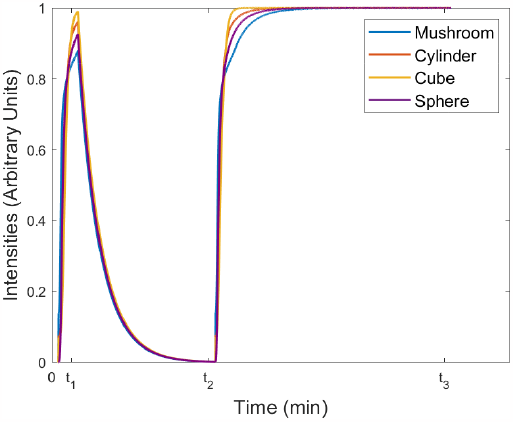
Total intensity of the potassium ions averaged over an entire biofilm as a function of time for different biofilm shapes spiked at their centres. The parameters were kept constant throughout the different geometries: the *K*^+^ ion diffusivity (*D*_*c*_) was 0.1 *µm*^2^*/s*, the total potassium ions added (*σ*) was 5 *×*10^9^, the firing threshold (*c*\*) was 1*×* 10^7^ and the decay rate (*k*_*s*_) was 5 *×*10^*−*3^. The first wavefront of the FDF stopped at *t*_1_ = 180 s and the second wavefront began (new hyperpolarization event) at *t*_2_ = 1440 s. The simulation stopped at *t*_3_ = 3600 s.

To understand the dependence of the squared radial displacement of the *K*^+^ wavefronts as a function of time for the different geometries, three parallel axes perpendicular to the y-axis were used (Figure III.17 (a)) where in Axis 1, the initial spike will take place instead of the centre for the cylinder with a defect. This allowed the wavefront propagation from three reference points surrounding a defect to be explored. This is shown in Figure III.18 for the different geometries. Different regimes are highlighted with trend-lines for different *γ* values. Furthermore, making these biofilms small, will also create an additional regime due to the size relative to the overall simulation space. This is shown in Figure III.19 for the smaller geometries.

**FIG. III.17.**
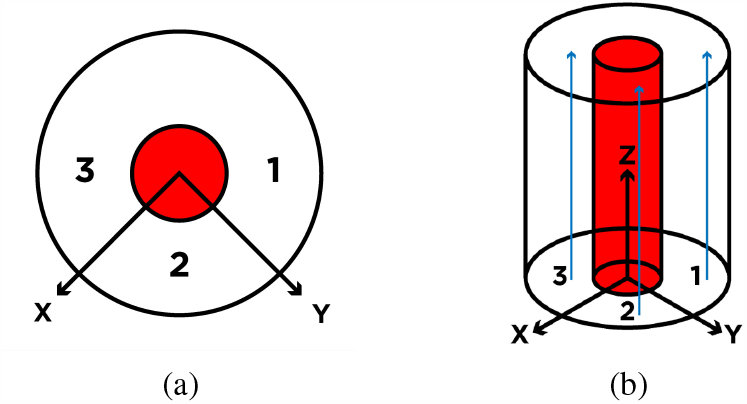
Schematic diagram of a cylindrical biofilm with a cylindrical defect shown in red used in the simulation. The bacteria are randomly placed in the cylinder except for the area shown in red. (a) Top view of the plane perpendicular to the y axis. (b) Side view of the plane parallel to the y axis. The blue arrows indicate the directions in which the wavefront propagation was quantified.

**FIG. III.18.**
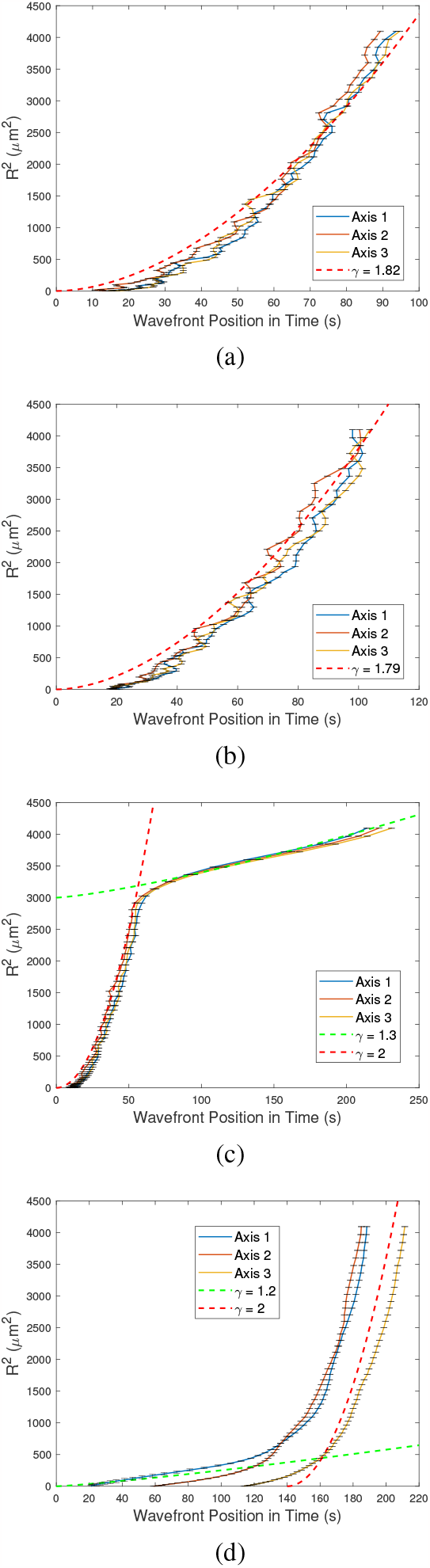
Mean square displacement, *R*^2^, of the *K*^+^ wavefront as a function of time for (a) a cylinder, (b) a cube, (c) a mushroom-like shape and (d) a cylinder with a cylindrical defect. The simulation parameters were kept constant for the different geometries. The *K*^+^ diffusivity (*D*_*c*_) was 0.1 *µm*^2^*/s*, the total *K*^+^ added (*σ*) was 5*×*10^9^, the firing threshold (*c*\*) was 1*×*10^7^ and the decay rate (*k*_*s*_) was 5*×*10^*−*3^. The trendlines for different *γ* values are shown for the different regimes. The error bars are the standard mean error on the wavefronts for 3 runs.

**FIG. III.19.**
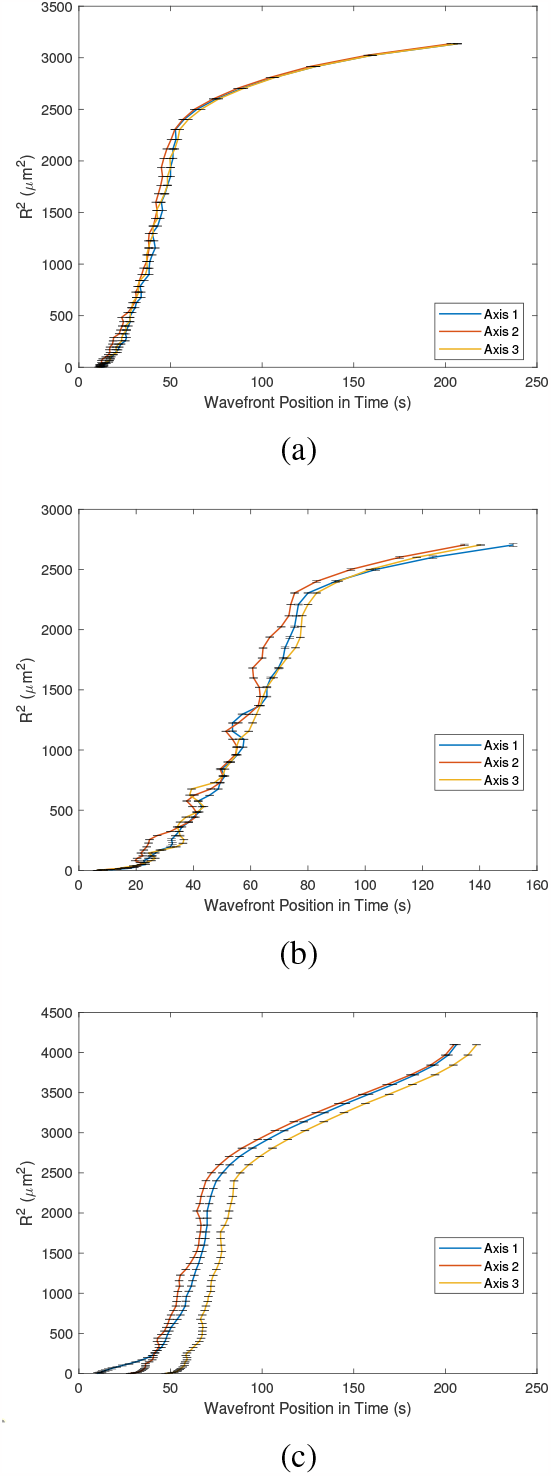
Mean square displacement, *R*^2^, of the *K*^+^ wavefront as a function of time for (a) a small cylinder, (b) a small cube, and (c) a small cylinder with a cylindrical defect. The simulation parameters were kept constant for the different geometries. The *K*^+^ diffusivity (*D*_*c*_) was 0.1 *µm*^2^*/s*, the total *K*^+^ added (*σ*) was 5*×*10^9^, the firing threshold (*c*\*) was 1 10^7^ and the decay rate (*k*_*s*_) was 5*×*10^*−*3^. The error bars are the standard mean error on the wavefronts for 3 runs.

### J. Release and Dormancy Times of *K*^+^ Ions by Bacteria

The kinetics of *K*^+^ ion release by the bacteria was investigated. Initially the concentration profile used for *K*^+^ release was similar to a delta function, Figure III.20 (a). The potassium ions are instantly released once the bacterium is above the threshold concentration (*c*\*) and the width of the pulse is limited by the time step of the simulation. More realistic behaviour is provided by adding a rise time for how long the cell will release the chemical and a dormancy time for when the cell will not fire even if it is above the firing threshold (similar to the refractory period of a neuron [7]), Figure III.20 (b). These release characteristics were used for a spherical biofilm and the *R*^2^ and total intensity were measured for different rise and dormancy times.

1, 5 and 10 s were chosen for both the rise and the dormancy times for the cells and the *R*^2^ of the *K*^+^ wavefront was plotted as a function of time, Figure III.21. A super-diffusive behaviour for the *K*^+^ wavefront dynamics was observed regardless of the rise and dormancy times. The curve shifts to later starting times with a higher dormancy time as expected.

**FIG. III.20.**
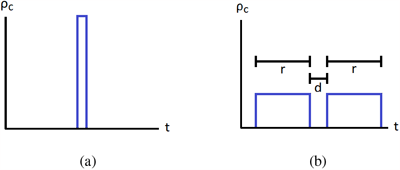
Concentration profile of potassium ions released from a single bacterium as a function of time (*t*). *ρ*_*c*_ is the potassium concentration density i.e. the *K*^+^ concentration per unit time. The area of the rectangles equals the total *K*^+^ concentration released. (a) The initial FDF model with instantaneous release and a finite time step. (b) A sustained release time (*r*) and a dormancy time (*d*) are shown.

**FIG. III.21.**
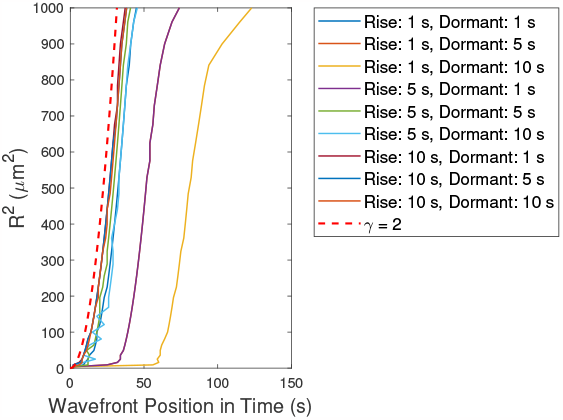
The mean square position (*R*^2^) of the *K*^+^ wavefront, *R*^2^, as a function of time for different rise and dormancy times from ABM simulations. The *K*^+^ diffusivity (*D*_*c*_) was 0.1 *µm*^2^*/s*, the total *K*^+^ added (*σ*) was 5*×* 10^9^, the firing threshold (*c*\*) was 1 10^7^ and the decay rate (*k*_*s*_) was 5*×* 10^*−*3^. The trendline is a power law with exponent *γ* = 2 (dashed line) showing ballistic scaling.

## IV DISCUSSION

The parameters used in the 3D ABM FDF model for *E. coli* were initially inspired by Blee *et. al*’s [16] simulations for *B. subtilis*. These parameters were then adapted based on more specific information from *E. coli* [6].

In Figure III.2 power law scaling occurs for the average *K*^+^ wavefront velocity on the diffusivity of the *K*^+^ ions. In Figure III.3, there is also a clear power law scaling for the instantaneous wavefront velocity on different *K*^+^ diffusivities and the velocity is larger for smaller radii. Furthermore, the power law exponents for the scaling of the instantaneous velocity are shown in Figure III.4, which depend on the radii. They fall in the expected range, 0.5-1, for a FDF model [7].

In Figure III.5, both simulation and experiment agree that the first peak is sharp and that the second wave plateaus. However, in the simulation, although the first peak is similar to experiments, there is a discrepancy between the rate at which the intensity increases (i.e. during hyperpolarisation of the cells) for the second rise. More detailed models of ion channel kinetics are needed to describe this e.g. Hodgkin-Huxley models for ion channel opening and models for cell to cell variability need to be considered. [7] [8]

The centrifugal *K*^+^ wavefront has a super-diffusive scaling exponent of *γ* of 1.21 for *R*^2^ on time, Figure III.6. For the centrifugal *K*^+^ wavefront, the critical radius is the minimum radius needed to nucleate a propagating wavefront. The centripetal *K*^+^ wavefront (Figure III.6) has a scaling exponent, *γ*, of 2.26, making it super-ballistic. Thus the nature of the dynamics changes as the wavefront collapses back into the biofilm.

The *K*^+^ wavefront velocity as a function of time for centrifugal waves is seen in Figure III.7 (a), where there is an inverse proportionality. Figure III.7 (b) shows in contrast that the velocity of centripetal waves increases with time. As the *K*^+^ wavefront is propagating towards the centre, the velocity increases monotonically. In Figure III.8 (a) the *K*^+^ centrifugal wavefront velocity varies below a threshold curvature value, also suggesting a critical value for the radius. Figure III.8 (b) shows for the centripetal *K*^+^ wavefronts the velocity increases and then saturates at high curvature which suggests that the larger curvatures contribute to the greater velocities. Both varieties of wavefront exhibit the plateau expected from the Eikonal approximation over a range of higher curvatures [7].

The *K*^+^ wavefront positions experience anomalous diffusion in the experimental data and are in agreement with the simulation. The centripetal wave is superballistic and centrifugal wave is subballistic and super diffusive. Furthermore, there is a slow decrease of the velocity for the centrifugal wavefront as a function of time in experiments [6], which looks similar to the model predictions in Figure III.7 (a). However, the centripetal velocity wavefront in Figure III.7 (b) is not in exact agreement with experiment. This suggests there is another mechanism unaccounted for in the model that limits its accuracy, such as heterogeneity of microclusters within the bacterial biofilms. It is suggested that *E. coli* biofilms have subpopulations of the cells with distinct phenotypes [3].

The scaling of the mean square wavefront position (*R*^2^) with the power law scaling factor *γ* on time for different *K*^+^ diffusion coefficients (Figure III.12) seems reasonable since the wavefront struggles to propagate for lower diffusion coefficients for the potassium ions and a more ballistic diffusion of the wavefront occurs.

In Figure III.13, higher values of the firing threshold (*c*\*) for *K*^+^ ion release makes the *K*^+^ wavefront behave more subdiffusively as a function of time. The *K*^+^ wavefront lingers more on each bacterium for high threshold values and therefore propagates slower.

Figure III.14 shows decreasing the diffusion coefficient of the *K*^+^ ions causes the critical radius of the propagating wavefronts to decrease. Above the maximum value of *K*^+^ diffusivity shown in the figure, the critical radius ceased to exist. Furthermore, at higher firing thresholds (*c*\*), there is a decrease of the critical radius.

The different biofilm geometries (sphere, cylinder, cube and mushroom) only cause a minor changes to the dynamics of the globally averaged potassium ion concentration and show the double hyperpolarization phenomenon observed in experiments, Figure III.16. The *R*^2^ values as a function of time in Figure III.18 show that for the cylinder and the cube, the system has similar super-diffusive dynamics to the spherical biofilm. However for the mushroom shape, the curve has a second regime where the superdiffusive behaviour changes to a diffusive or subdiffusive-like behaviour at short time scales. The beginning of the three curves of the *R*^2^ as a function of time for the cylinder with a defect (Figure III.18 d) all start as expected with axis 1 going first (it is closest to the point of stimulation) and axis 2 starting after a delay time and axis 3 starting after a longer delay. The behaviour quickly changes from a subdiffusive to a superdiffusive behaviour, suggesting that the waves propagate differently once the whole geometry along the axis is activated.

Interestingly, if the biofilm geometries are made smaller, a subdiffusive characteristic appears in all the geometries, similar to the mushroom-like shape. This suggests that the superdiffusive transport of *K*^+^ wavefronts at long times combines with the subdiffusive behaviour at short times in biofilm geometries that have lower dimensionalities.

When introducing release and dormancy times for the firing of the potassium ions, the *K*^+^ wavefronts are not very different in terms of the superdiffusion of the wavefront position squared (*R*^2^) as a function of time. However, the *K*^+^ wavefront starts with different delay times, Figure III.21. This suggests that the transport of the *K*^+^ wavefront does not depend on the release and delay times and will be relatively robust to the opening and closing rates of the ion channels. Furthermore, the intensity profile is rescaled in amplitude by the release and dormancy times without changes to the shape. This suggests that the release and dormancy times only influence the maximum *K*^+^ wavefront amplitude and the process is robust to minor variations in these parameters.

Anomalous diffusion of wavefronts in reaction-diffusion systems has not been studied extensively in the literature from the perspective of simulations [16] or experiments [6]. Analytic solutions to classical reaction-diffusion equations tend to predict diffusive (*R*^2^*∼t*) or ballistic (*R*^2^*∼t*^2^) scaling of wavefront transport, but it is an inconvenient truth that the majority of real systems probably have intermediate scaling of wavefront position with time [27]. For the anomalous motion of single particles, standard models invoke non-Markovian effects (e.g. continuous time random walks or fractional Brownian motion) [27]. No non-Markovian effects were explicitly included in our simulations (classical diffusion was modelled for the ions and ion release from bacteria is assumed fast on the time scale of the simulations e.g. there are no fat tails in the probability distributions for ion release), so the physical origin of the anomalous wavefront dynamics is currently unclear and requires further research. At long time scales (*t→ ∞*) and long distances from a biofilm (*R→ ∞*) the concentration profile of *K*^+^ ions must resemble diffusion from a point source, so diffusive kinetics of ion transport are expected in the limiting regimes of long time and long distance scales. Furthermore, wavefront dynamics will terminate once all the bacteria have released their potassium e.g. at the edge of the biofilm for centripetal wavefronts. Thus the anomalous kinetics of wavefronts will only occur over an intermediate range of time scales, but this is expected to be biologically relevant and corresponds to an experimentally accessible time window [16] [6].

It would be interesting to segment confocal microscopy images of a real biofilm and then predict the wave fronts on real biofilm communities based on the FDF ABM. Percolation transitions for the propagation of electrical wavefronts are possible in mixed species biofilm, where one species is more susceptible to hyperpolarization. Different types of electrical signal quenching are also possible analogous to the phenomenon of *quorum quenching* which is observed biologically [32] and could be explored with ABM.

The model could also be further developed by adding different species in the model. That is, by having agents with different characteristics, such as the threshold for firing and spatial positions, could provide a more realistic interpretation of common biofilms which contain many species of bacteria [3].

It is possible that the electrical response of many bacterial cells is anisotropic e.g. gap junctions have been discovered in marine bacteria [33]. It would be interesting to model the effect of electrical anisotropy (e.g. to place more ion channels on the axis of elongated cells) on the propagation of *K*^+^ wavefronts.

Future simulations can be developed that include the full Hodgkin-Huxley equations for the neuronal-like activity of the bacteria [7]. The current fire-diffuse-fire model provides a minimal model to describe a series of phenomena in light-stressed *E. coli* biofilms e.g. the anomalous transport of wavefronts and the critical radius for wavefront propagation. However, additional experiments using electrical stimulation of *E. coli* biofilms demonstrate some non-linear aspects of the hyperpolarization phenomena that will require a non-linear Hodgkin-Huxley model for their explanation e.g. hysteresis in *K*^+^ release. Electrical impedance spectroscopy measurements with *E. coli* biofilms also are providing additional information on the activity of ion channels that need to be included in agent based models e.g. neuromorphic negative capacitances due to *K*^+^ ion channels [34] [35].

## V. CONCLUSIONS

Fire-diffuse-fire agent based modelling provides a satisfactory description of many of the emergent phenomena observed in the electrical signalling of bacterial biofilms e.g. when *E. coli* biofilms experience stress to blue light. Anomalous dynamics of the wavefronts are well described with a relatively simple fire-diffuse-fire mechanism calculated numerically with agent based modelling. A critical radius for *K*^+^ wavefront propagation is an emergent property of the simulations and is observed in experiments. The dependence of *K*^+^ wavefront propagation on the biofilm geometry and defects in the biofilms was investigated and it could motivate future experiments e.g. to observe geometry dependent transitions between superdiffusive and subdiffusive scaling of the *K*^+^ wavefronts.

## Supporting information

Supplementary Information Tables and Video Description

Cube Potassium Propagation Video

Cylinder With Defect Potassium Propagation Video

Mushroom Like Shape Potassium Propagation Video

Cylinder Potassium Propagation Video

Sphere Potassium Propagation Video

## ACKNOWLEDGMENTS

We thank Johanna Blee, Elyece Malnati, Kiera Moreno, Emily Gillot and Harry Lipscomb for their contributions, as well as Jose Carlos Bre ña for his help with image design. Further, Ovidio Cesar and Thiago Militino for their assistance in Java coding.

