## Supplementary Information Tables and Video Description for "Electrical Signalling in Three Dimensional Bacterial Biofilms Using an Agent Based Fire-Diffuse-Fire Model"

(Dated: November 15, 2023)

### I. SUPPLEMENTARY TABLES

General parameters for the simulation of electrical signalling in three dimensional *E. coli* biofilms are shown in Table I. The parameters for Blee at al's model [1] for *B. subtilis* biofilms in 2D for comparison are shown in Table II.

| Model Parameter | Value | Units |
|---|---|---|
| Simulation Time Step ($dt$) | 1 | $s$ |
| Grid Size | $64 \times 64 \times 64$ | $\mu m^3$ |
| First Wave Stoppage Time | 180 | $s$ |
| Second Wave Beginning Time | 1440 | $s$ |
| Diffusivity ($D_c$) | 0.1 | $\mu m^2/s$ |
| Decay Rate ($k_s$) | $5\times10^{-3}$ | $molecules/s$ |
| Quantity of Potassium Added Per Spike ($\sigma$) | $5\times10^{9}$ | $molecules$ |
| Total Number of Bacterial Agents | 4500 for Mushroom Shape<br>6000 otherwise | – |
| Fire-Diffuse-Fire Threshold ($c^*$) | $1\times10^{7}$ | $molecules$ |

TABLE I. Supplementary table for the simulation parameters used for the FDF ABM of 3D *E. coli* biofilms. These were the default parameters, but they were modified to understand functional dependencies in the main text.

* Biological Physics, Department of Physics and Astronomy, University of Manchester, Oxford Rd., Manchester, M13 9PL, UK.; Division of Infection, Lydia Becker Institute of Immunology and Inflammation, School of Biological Sciences, University of Manchester, Oxford Rd., Manchester, M13 9PT, UK.

† Division of Evolution, Infection and Genomics, School of Biological Sciences, Faculty of Biology, Medicine and Health University of Manchester, Manchester, M13 9PT, UK.

‡ Division of Infection, Lydia Becker Institute of Immunology and Inflammation, School of Biological Sciences, University of Manchester, Oxford Rd., Manchester, M13 9PT, UK.;

§ Biological Physics, Department of Physics and Astronomy, University of Manchester, Oxford Rd., Manchester, M13 9PL, UK.; Photon Science Institute, Alan Turing Building, Oxford Rd, Manchester M13 9PY, UK;

| Model Parameter | Value | Units |
|---|---|---|
| Simulation Time Step (dt) | 21 | $s$ |
| Pixel Size | 0.1 | $\mu m$ |
| Signal Grid Length | 20 | $pixel(px)$ |
| Signal Grid Cell Size | 20 | $px^2$ |
| Diffusivity | 0.4 | $molecules\ cell\ grid/dt$ |
| Decay Rate | $7{\times}10^{-2}$ | $molecules/dt$ |

TABLE II. Supplementary table for the simulation parameters used for Blee et al's ABM simulations for *B. subtilis* in 2D [1]. As the models differ, there were values which were not present in the current model compared to Blee et al's model and vice versa. The most relevant parameters for comparison to our model were included in this table.

### II. SUPPLEMENTARY VIDEOS

The supplementary videos show how the different geometries influence the propagation of the potassium field. The number at the top left of the videos represents seconds. The video was sped up so that 3600 seconds was shown over 40 seconds for all geometries. Outlines of the shapes formed from the spatial locations of the bacteria in the biofilms were provided in each video.

The videos show the spiking in the bottom of the biofilm for a mushroom shape, a cylinder and a cube. A video also shows the spiking of a spherical biofilm in the centre. These can be appreciated in Figure III.16 and in the case of the sphere, in subfigure b) in Figure III.5. The sphere size was 32 $\mu m$ in radius i.e. the diameter fills the whole workspace. The mushroom was a complex shape comprised of a hemisphere and a one-sheet hyperboloid placed below this hemisphere [2]. The hemisphere has a radius of 19.36 $\mu m$ and the one-sheet hyperboloid has a radius of 260.27 $\mu m$. The cylinder had a height of 64 $\mu m$ and a radius of 25.6 $\mu m$. The cube was slightly smaller than the whole workspace to allow a better visualisation with length of 48 $\mu m$

Another video shows the propagation of the potassium field of a cylinder with a defect in which the spike occurs at the bottom of the cylinder along axis 1. The parameters used for the simulation are the same for the cylinder with the defect as the other geometries (Table I). Furthermore, the radius of the defect was a factor of 0.7 of the total radius, which was 32 $\mu m$, with a height of 64 $\mu m$.

[1] J. A. Blee, I. S. Roberts, and T. A. Waigh, Spatial propagation of electrical signals in circular biofilms: A combined experimental and agent-based fire-diffuse-fire study, Physical Review E **100**, 052401 (2019).

[2] B. R. Martinand S. G., *Mathematics for physicists* (John Wiley  Sons, 2015).
